# Beyond Gloger’s rule: multiple biogeographic drivers of dark and red colouration in ants

**DOI:** 10.64898/2026.03.16.712184

**Authors:** Tristan Klaftenberger, Brian L. Fisher, Eddie Pérochon, Cleo Bertelsmeier, Sébastien Ollier

**Author notes:** Equal contribution (Cleo Bertelsmeier, Sébastien Ollier).

## Abstract

Colour is a key trait involved in camouflage, physiological protection and thermoregulation. Yet environmental drivers of colour variation remain poorly understood at large spatial scales. Gloger’s rule predicts animals should be darker in warmer and wetter climates, and in a complex version, redder in warmer and drier climates. Here, we present the first test of the complex Gloger’s rule in insects using 34,331 images of 10,400 ant species across 586 assemblages worldwide. We decompose species mean colouration into two orthogonal axes, linked to darkness and redness. Assemblages were darker under high UV-B radiation and low dry-season precipitation, consistent with UV protection and desiccation resistance via melanisation. Canopy height increased both axes, suggesting camouflage. In contrast, higher mean temperature of the warmest quarter increased redness, as predicted by the complex Gloger’s rule. Ant colouration cannot be explained by one macroecological rule but reflects environmental drivers acting independently on darkness and redness.

## 1 Introduction

Animal colouration plays a central role in survival and reproductive success, mediating a wide range of functions including camouflage, physiological protection, and thermal regulation (Cuthill *et al*. 2017). However, colour is often shaped by multiple selective pressures that may act in opposing directions (Cuthill *et al*. 2017). Consequently, identifying the dominant drivers of colour variation and disentangling their relative contributions remains challenging, particularly at broad spatial scales.

One of the earliest attempts to describe large-scale geographic patterns of animal colouration is Gloger’s rule (Rensch 1929), which in its simplest formulation, predicts that animals should exhibit darker colouration in warm and humid environments, driven by increased melanin deposition (Delhey 2019). However, Rensch’s original formulation (generally referred to as the ‘complex version of Gloger’s rule’) already distinguished between different components of melanin-based colouration and their potentially distinct responses to environmental gradients (Rensch 1929). This complex version explicitly distinguishes between eumelanin-based darkness and pheomelanin-based redness, and their predicted responses to temperature and humidity gradients (Delhey 2019). Along temperature gradients, both pigmentation types are predicted to increase from cold to warm environments. Contrarily, their responses diverge along humidity gradients: eumelanin deposition is predicted to increase from dry to humid environments, while pheomelanin deposition is predicted to increase under dry conditions (Fig. 1).

**Figure 1.**
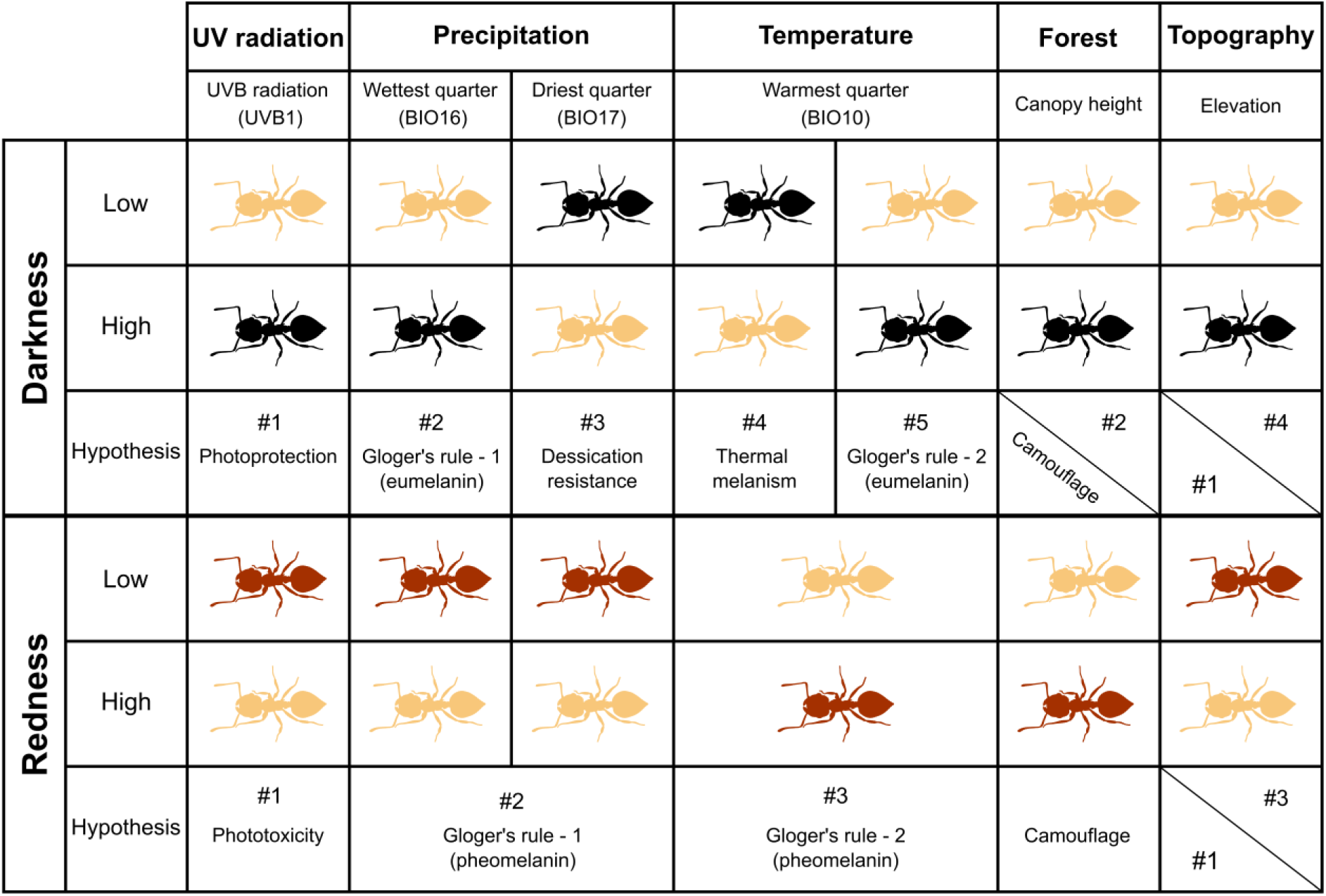
Conceptual framework linking environmental gradients to melanin-based colour variation. Predicted responses of eumelanin-based darkness (upper panel) and pheomelanin-based redness (lower panel) along major environmental gradients. Columns represent broad classes of environmental factors, while subheadings indicate the specific variables used in this study as proxies to best capture each hypothesis. Ant silhouettes illustrate the expected direction of colour change between low and high values of each variable.

The mechanisms underlying these predictions are debated and likely non-exclusive (Delhey 2019). Increased eumelanin-based dark pigmentation in humid environments has often been interpreted as a physiological response to elevated pathogen pressure, as eumelanin may enhance resistance to parasites and pathogens though pleotropic effects in vertebrates (Ducrest *et al*. 2008) or for example via encapsulation in insects (Dubovskiy *et al*. 2013; Wittkopp & Beldade 2009). Pathogen loads tend to be higher under humid conditions associated with forested habitats. However, forests are also characterised by shadier environments, which may favour darker colouration through improved camouflage (Delhey 2017). Consequently, both pathogen-mediated selection and camouflage predict darker animals in forested environments (Fig. 1). Eumelanin deposition is also predicted in warmer environments, but without a clearly established mechanism. One possible explanation is that warmer regions are often associated with higher levels of ultraviolet radiation, and increased melanisation may provide protection against UV-damage (Clusella-Trullas & Nielsen 2020; Delhey 2019; Roulin 2014). Under this photoprotection hypothesis, darker colouration is expected to increase with UV exposure rather than temperature *per se* (Fig. 1).

Importantly, the complex Gloger’s rule also predicts increased pheomelanin-based redness in warm and dry environments, but with less theoretical support (Delhey 2019; Rensch 1929). One potential mechanism is that pheomelanin can generate oxidative damage when exposed to ultraviolet radiation (phototoxicity hypothesis) (Ito *et al*. 2018). Under this hypothesis, increased UV exposure is expected to constrain or reduce pheomelanin-based redness, independently of temperature or humidity (Fig. 1). Another possibility is that forest environments may select for increased redness through camouflage. Forests are characterised by low luminance and light spectra dominated by green wavelengths, as vegetation absorbs red and blue light (Endler 1993). Under these conditions, reddish colouration may be less perceptible, particularly when combined with increased darkness, favouring darker brownish phenotypes produced by the joint expression of eumelanin and pheomelanin. Similar crypsis-related mechanisms have been documented in tropical birds, where species inhabiting the forest understorey exhibit browner and redder plumage compared with canopy-dwelling species, consistent with improved background matching in dim forest light environments (Gomez & Thery 2007; Gomez & Théry 2004).

Despite its prominence as one of the top seven major macroecological rules (Tian & Benton 2020), empirical evidence for Gloger’s rule is often partial, geographically or taxonomically restricted and sometimes even contradictory or inconsistent (Delhey 2019). Support at the global scale has been reported for some taxa, notably passerine birds (Delhey *et al*. 2019), which has led to the widespread assumption that Gloger’s rule is one of the best documented macroecological patterns. First, most studies have however only tested if either humidity or precipitation alone were linked to variations in darkness, and have not assessed both drivers simultaneously (Delhey 2019). Second, only a handful of studies have explicitly examined variations in pheomelanin-based colouration jointly with variations in eumelanin-based darkness, consistent with predictions of the complex Gloger’s rule. These studies have focused exclusively on endothermic vertebrates, including domestic pigs (Newell *et al*. 2021), rodents and lagomorphs (Cerezer *et al*. 2024, 2025), mammals more broadly (Howell & Caro 2025), or birds (Marcondes *et al*. 2020, 2021) and have yielded mixed support for the complex Gloger’s rule. To date, no equivalent joint test of darkness and redness predictions has been conducted in ectotherms.

This evident knowledge gap may be due to the complexity of addressing this question, particularly in ectotherms. For this group of animals, two additional hypotheses may be particularly important in explaining variations in colour, complicating things further. First, the desiccation-resistance hypothesis (Fig. 1) links darker colouration with dry environments because melanin reduces cuticular permeability and water loss in insects (Clusella-Trullas & Nielsen 2020; Parkash *et al*. 2008a, 2009b; Ramniwas *et al*. 2012). Second, Bogert’s rule, also known as the thermal melanism hypothesis (Bogert 1949; Clusella Trullas *et al*. 2007), predicts darker colouration in colder environments favouring heat absorption, and lighter colouration in hot environments reducing overheating (Fig. 1). Finally, elevational gradients may combine both mechanisms (Bishop *et al*. 2016), increased UV exposure and lower temperatures are expected to select for darkness (thermal melanism, UV protection) and reduced pheomelanin-based redness at higher elevations (phototoxicity) (Fig. 1).

Taken together, these mechanisms generate several alternative predictions linking colour variation to environmental gradients (Fig. 1). Specifically, we tested four non-exclusive hypotheses: (1) predictions derived from the complex formulation of Gloger’s rule, expecting eumelanin-based darkness to increase with temperature and humidity, and pheomelanin-based redness to increase with temperature but decrease with humidity; (2) the photoprotection hypothesis, predicting darker colouration under higher ultraviolet radiation; (3) the desiccation-resistance hypothesis, predicting darker colouration in dry environments; and (4) habitat-mediated camouflage, predicting darker and redder colouration in forested environments where background matching may be favoured.

Here, our goal was to disentangle the relative contributions of these different mechanisms to large-scale variation in ant colouration. By jointly considering variables that best represent each hypothesis, we assess how multiple, potentially non-exclusive mechanisms contribute to large-scale variation in colouration. To achieve this, we chose ants (Hymenoptera; Formicidae) as a study system because they are among the most ecologically dominant insect lineages and occur across nearly all terrestrial ecosystems. Moreover, the >14,000 described species of ants are estimated to collectively exceed the combined biomass of wild mammals and birds (Hölldobler & Wilson 1990; Schultheiss *et al*. 2022). Previous work has shown that ant colour variation is dominated by a pale–dark gradient, while most species are restricted to red hues (Idec *et al*. 2023). This combination makes ants particularly well-suited for jointly examining geographic variation in both darkness and redness. To do that, we used a global dataset of ant occurrence records (Kass *et al*. 2022) to compute assemblage-level mean values of darkness and redness, enabling us to test the full set of hypotheses at a global scale (Fig. 1). In addition, we describe broad geographic patterns in assemblage colour composition, including latitudinal trends, to provide a global overview of colour variation before examining environmental drivers.

## 2 Methods

### 2.1 Colour data acquisition and processing

Ant body colour was quantified using standardized specimen images from AntWeb (www.antweb.org), a global repository of ant images and associated metadata. All photographs were taken following a standardized imaging protocol, allowing direct comparisons across species and geographic regions (Idec *et al*. 2023).

We analysed profile-view images of worker specimens only, using a total of 34,331 images representing 10,400 species. For each specimen, a single observer selected one pixel from the mesosoma, avoiding areas affected by glare or underexposure. Colour values were extracted in the HSV colour space, where hue, saturation and value describe colour type, purity and brightness, respectively.

The mesosoma was selected because its colouration is expected to be closely linked to cuticular melanisation and may therefore better reflect environmental conditions. In contrast, head colour may partly reflect structural reinforcement: strong mandibular muscles impose mechanical stresses on the head capsule, which can promote local cuticle thickening and sclerotisation/melanisation, potentially affecting apparent pigmentation (Andersen 2012; Fraenkel & Rudall 1940; Matte *et al*. 2025). Gaster colour can also vary with physogastry and specimen preservation.

It was not possible to account for intraspecific colour variation which has been reported in ants (Idec et al. 2023; Skaldina & Sorvari 2017a) because sampling across species was highly uneven. Approximately one third of species (∼33%) were represented by a single specimen with valid colour data (mean number of specimens per species = 3.3; median = 2, maximum = 164). As a result, intraspecific variation could not be reliably quantified for most species and was therefore not analysed. Instead, colour values were averaged at the species level using arithmetic means of HSV components. Species-level mean colour values are reported for all species included in the study. In addition, specimen-level colour data for a randomly selected set of 50 species are available in the data repository associated with this study, enabling independent verification of the species-level averaging procedure.

To summarise interspecific colour variation, we conducted a mean-centred principal component analysis (PCA) on seven colour descriptors derived from three colour spaces (HSV, CIE L*u*v*, and RGB), using the package ade4 (Dray & Dufour 2007) (v1.7.22). Hue was excluded from the HSV system because it is a circular variable scaled between 0 and 1, with extreme values representing adjacent colours on the colour wheel. Treating hue as linear would therefore violate the Euclidean distance assumptions underlying PCA. Because green is almost absent in ants, and lighter-coloured ants generally appear yellow, the red and green channels of the RGB system were combined into a single yellow component (R + G). Species scores on the first two principal axes were retained for subsequent analyses.

### 2.2 Ecoregional ant assemblages

To examine biogeographical drivers of variation in mean colour at the assemblage level, we defined ant assemblages as sets of species occurring within similar macroenvironmental conditions. Assemblages were delineated using the world’s 846 terrestrial ecoregions, which represent ecosystems of regional extent (Dinerstein *et al*. 2017) and are widely used in large-scale biogeographical analyses (Vynne *et al*. 2022; Yu *et al*. 2019), including studies of large-scale patterns of bird colouration (Cooney *et al*. 2022) or ant societies (Pérochon *et al*. 2026).

Species composition within each ecoregion was characterised using a global database of ant occurrences comprising 1,479,293 georeferenced records from 14,328 species (Kass *et al*. 2022). Occurrence points were spatially intersected with ecoregion polygons using the sf package (v1.0.16), after verifying polygon validity. A total of 18,116 records could not be assigned to any ecoregion and were excluded, resulting in a final dataset of 1,461,181 occurrences representing 14,317 species.

Global ant occurrence data are affected by strong spatial heterogeneity in sampling effort, which can generate spurious biogeographical patterns. To address this issue, we relied on a recently published evaluation of sampling completeness in global ant occurrence records, in which political regions were assessed by comparing georeferenced occurrences with species presence lists (Pérochon et al. 2026). In this assessment, regions were classified as well-sampled based on a combination of complementary criteria capturing spatial coverage, taxonomic completeness, phylogenetic representativeness, and compositional similarity between occurrence data and regional species lists. Political regions meeting all criteria were intersected with terrestrial ecoregions, and ecoregions were considered well-sampled when at least half of their area overlapped with well-sampled political regions. This procedure identified 661 terrestrial ecoregions as well-sampled. Of these, 28 contained no ant occurrences, leaving 633 ecoregions.

Among these ecoregions, a small number still contained very few species despite being classified as well-sampled. This may reflect remaining data gaps but could also correspond to regions that naturally harbour very low ant species richness. To reduce the influence of such extreme cases, we applied a 5% richness filter, excluding the lowest 5% of ecoregions ranked by species richness. This procedure removed 35 ecoregions, leaving 598 ecoregions for subsequent analyses. Because several ecoregions shared the same species richness at the threshold value, all tied cases were removed, resulting in a slightly larger exclusion than the nominal 5%.

Finally, using the list of species recorded in each ecoregion and their PCA1 and PCA2 scores, we calculated the mean score of all species present in each ecoregion, representing the average position of assemblages along the two main axes of colour variation.

### 2.3 Explanatory variables

To test all hypotheses, we selected a set of six bioclimatic and environmental predictors that capture the main climatic, radiative, and habitat-structural dimensions relevant to colour variation, while remaining sufficiently independent to be analysed jointly (Fig. S1).

Climatic variables were extracted from the WorldClim v2.1 database (Fick & Hijmans 2017) at a spatial resolution of 30 arc seconds. To test temperature-related hypotheses, we used the mean temperature of the warmest quarter (BIO10), which captures thermal conditions during periods of peak heat and avoids averaging over seasons when insects are inactive in many regions.

To test humidity-related predictions, we specifically selected two precipitation variables that allow us to distinguish between Gloger’s rule (precipitation of the wettest quarter (BIO16)) and the desiccation-resistance hypothesis (precipitation of the driest quarter (BIO17)) in the context of darkness variation. For consistency, we used the same variables when testing predictions for redness. These variables capture climatic extremes during which selection associated with high moisture availability or with water limitation is expected to be the strongest.

To test hypotheses related to UV radiation, including photoprotection and phototoxicity, we used UVB1 (mean annual UV-B irradiation) from the glUV radiation dataset for macroecological studies (Beckmann *et al*. 2014), which has been widely applied in global analyses of animal pigmentation, including previous macroecological studies of ant colouration (Bishop *et al*. 2016; Idec *et al*. 2023; Klunk *et al*. 2022).

We also integrated two environmental descriptors capturing topographic and habitat-structural variation. Elevation was extracted from WorldClim at 30 arc-second resolution (Fick & Hijmans 2017), and has previously been shown to be associated with darker ant assemblages (Bishop *et al*. 2016). Forest canopy height was used as a proxy for habitat structural complexity where lower canopy height reflects more open habitats and was already used to explain large scale variation of colouration (Cerezer *et al*. 2025; Goldenberg *et al*. 2025). Canopy height data were obtained from a global dataset available at 10 m resolution (Lang *et al*. 2023) and were downscaled to 30 arc seconds before extraction to match the resolution of other predictors.

For each predictor, ecoregion-level values were computed as the mean of 5,000 points randomly sampled within each ecoregion, ensuring a robust representation of within-ecoregion environmental conditions.

Species richness was quantified as the total number of ant species recorded within each ecoregion. Species occurrence data were obtained from a global ant occurrence database (see “Ecoregional ant assemblages”). To reduce skewness and limit the influence of extremely species-rich ecoregions, species richness was log-transformed before analyses.

Because ecoregions differ in the evolutionary lineages they contain, regions with similar phylogenetic composition may show similar trait distributions independent of environmental conditions. This is expected because the geographic distribution of ant diversity is strongly structured by macroevolutionary and biogeographic processes (Doré *et al*. 2025; Economo *et al*. 2018), and phylogenetic composition is known to influence functional trait structure among assemblages (Arnan *et al*. 2015; Cavender-Bares *et al*. 2009; Webb *et al*. 2002). To account for this, we quantified phylogenetic similarity among ecoregions. The underlying phylogenetic similarity matrix was retrieved from a previous study (Pérochon *et al*. 2026). This matrix quantifies the similarity in phylogenetic composition among ecoregions based on shared evolutionary lineages derived from a species-level phylogeny of ants (Economo *et al*. 2018). Because phylogenetic similarity values were not available for all ecoregions retained, 12 ecoregions were excluded, resulting in a final dataset of 586 ecoregions used in all subsequent analyses. We summarised this high-dimensional matrix using non-metric multidimensional scaling (NMDS), which provides a low-dimensional representation of pairwise similarities while preserving their rank structure. NMDS was performed using the vegan package (Oksanen *et al*. 2024; v2.6.6.1), and the two first dimensions were retained. In this ordination, ecoregions that are close in NMDS space have more similar phylogenetic structure (Fig. S5). The first NMDS axis was strongly correlated with species richness (r = −0.81; Fig. S1) and was therefore excluded from subsequent analyses. We retained NMDS axis 2 as a covariate representing phylogenetic similarity among assemblages while remaining largely independent of species richness (Fig. S1).

All analyses were conducted at the ecoregion level, with each ecoregion representing a distinct ant assemblage. Assemblage-level mean values of PCA1 (darkness axis) and PCA2 (redness axis) were used as response variables in separate models. All explanatory variables were standardised (mean = 0, SD = 1) prior to analyses to allow direct comparison of effect sizes. Species richness and ecoregion area were log-transformed before standardisation. To avoid collinearity, we examined pairwise correlations among all candidate predictors and retained only variables with absolute Pearson correlations below r > 0.7 (Fig. S1), a commonly used rule-of-thumb threshold to limit multicollinearity in ecological models (Dormann *et al*. 2013). The final set of predictors included species richness, phylogenetic similarity (NMDS2), ecoregion area, climatic variables (BIO10, BIO16, BIO17), UV-B radiation (UVB1), elevation, and canopy height. The global distribution of phylogenetic similarity among ecoregions further illustrates the strong spatial structuring of evolutionary lineages (Fig. S5).

### 2.4 Analyses

All analyses were conducted in R v. 4.4.1 (R Core Team 2024).

We analysed variation in assemblage-level colour using spatial linear mixed models fitted with the sdmTMB package (Anderson *et al*. 2025). This approach allows us to model spatial autocorrelation among ecoregions while estimating the independent effects of environmental predictors on colour variation. Spatial dependence among observations was modelled using a latent Gaussian random field implemented via the stochastic partial differential equation (SPDE) approach. This framework approximates a continuous spatial field with Matérn covariance on a triangulated mesh (a spatial network of connected triangles used to discretise the study area) (Lindgren *et al*. 2011). Triangulated spatial meshes were based on the geographic centroids of ecoregions. Mesh resolution was controlled by a cutoff distance parameter determining the minimum allowed distance between mesh vertices. Fixed effects included all standardised predictors described above, while the spatial random field captured residual spatial structure not explained by environmental variables. Models were fitted assuming a Student’s t observation distribution to accommodate heavy-tailed residuals. We systematically evaluated candidate meshes used to define the spatial structure of the models across cutoff distances ranging from 100 to 2000 km, using increments of 10 km. For each cutoff value, we fitted an otherwise identical spatial model and compared model support using the Akaike Information Criterion (AIC). Because AIC values are only meaningful for well-converged models, we retained only models that converged successfully and passed standard diagnostic checks using the sanity function from the sdmTMB package. The final mesh cutoff was selected as the cutoff yielding the lowest AIC among the convergent models, resulting in a cutoff of 500 km for the model fitted to assemblage-level mean PCA1 scores (darkness axis) and 400 km for the model fitted to assemblage-level mean PCA2 scores (redness axis).

Model fit was evaluated using conditional residual diagnostics, inspection of fitted versus observed values, and Moran’s I tests on MCMC-based quantile residuals to confirm the absence of residual spatial autocorrelation (Fig. S2; Fig. S3). Because p-values are not directly interpretable for spatial mixed models fitted via maximum likelihood, inference was based on the magnitude and direction of standardised regression coefficients and their confidence intervals. We report confidence intervals at multiple levels (90%, 95%, 99%, and 99.9%; Table S1 and Table S2).

To visualise broad latitudinal patterns in assemblage colour, we fitted generalized additive models (GAMs) relating assemblage-level mean PCA1 and PCA2 scores to the north–south position of ecoregion centroids in a Mollweide equal-area projection.

## 3 Results

### 3.1 Major axes of colour variation in ants

Principal component analysis of species mean colour across 10,400 ant species revealed two major axes of colour variation (Fig. 2). The first principal component (PCA1) explained 73% of the total variance and captured a pronounced pale–dark gradient. Low PCA1 scores corresponded to paler species, characterised by high values of luminance (L* in CIELUV) and value (V in HSV), whereas higher PCA1 scores corresponded to darker species, with lower values of these variables. In addition, paler species (low PCA1 scores) exhibited stronger yellow components, as indicated by higher values of the yellow channel (R + G in RGB) and the U coordinate in CIELUV.

**Figure 2.**
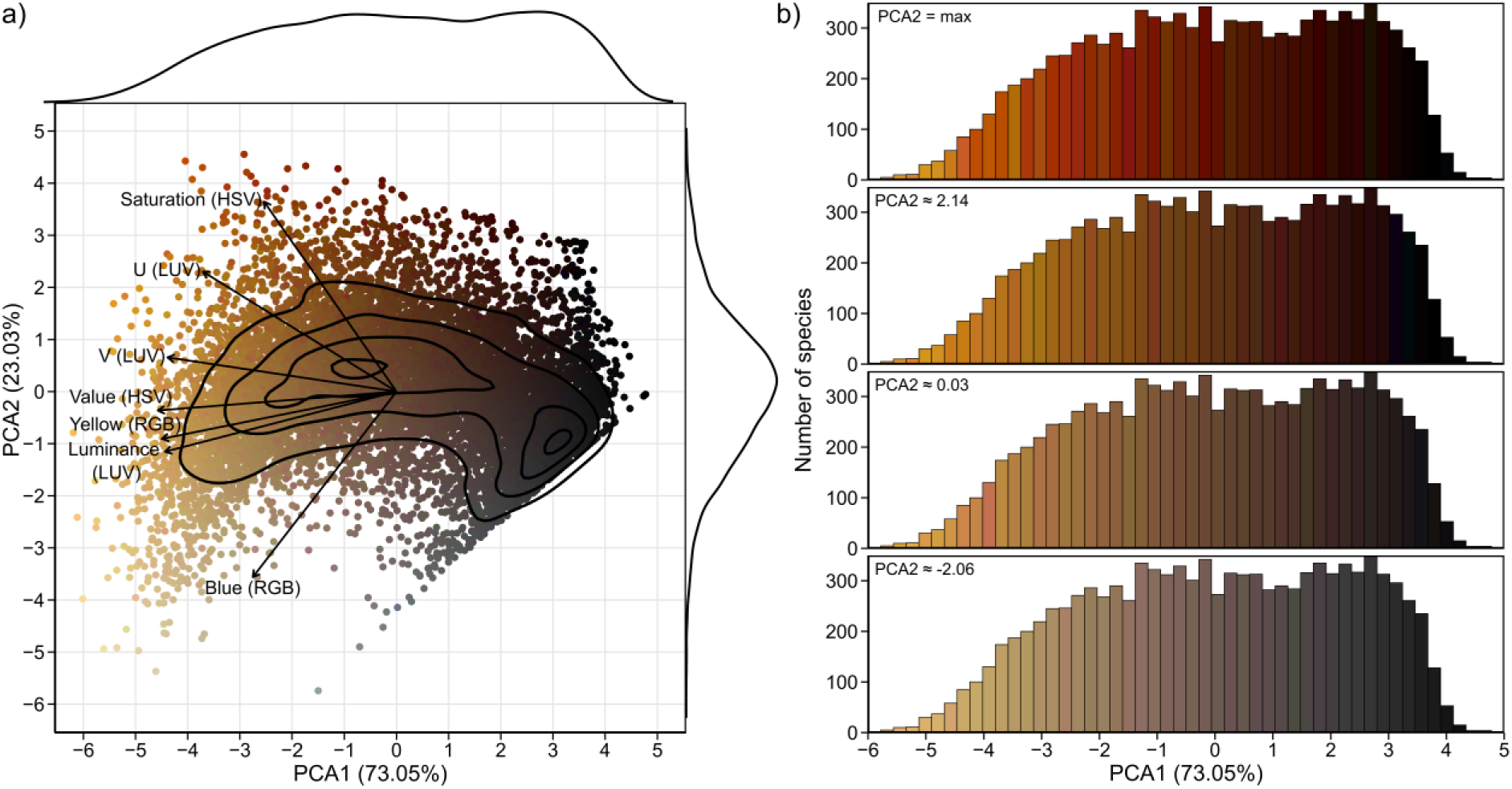
Major axes of colour variation across ant species. (a) Principal component analysis (PCA) summarising species’ average colour variation across 10,400 ant species using seven colour variables measured in three colour spaces (RGB, HSV excluding hue, and CIELUV). Species scores are shown on the first two components (PCA1 = 73.05% and PCA2 = 23.03% of total variance), with points coloured by each species’ average colour. Black contour lines show bivariate kernel density isoclines based on normalised density (ndensity = 0.25, 0.50, 0.75 and 0.95). Arrows indicate variables. Marginal density curves (top and right) show the distributions of species scores along PCA1 and PCA2. (b) Species-score distributions along PCA1. To visualise how colours vary along PCA1 at different positions on PCA2, bars are coloured using the average colour of species (points in panel a) whose PCA2 score is closest to the target PCA2 value within each PCA1 bin. From top to bottom, target PCA2 values correspond to the maximum observed value (PCA2 = max; the colour of the point with the highest PCA2 score in each bin), the 95% quantile (PCA2 ≈ 2.14), the median (50%; PCA2 ≈ 0.03), and the 5% quantile (PCA2 ≈ -2.06).

The second principal component (PCA2) explained 23% of the variance and described variation along a redness axis. Higher PCA2 scores were associated with redder and more saturated colours, characterised by higher values of the U coordinate in CIELUV and saturation in HSV. Conversely, lower PCA2 scores corresponded to less saturated colours with greater contributions from the blue channel in RGB and lower values of U and saturation. Visualisation of PCA1 score distributions with bars coloured by species colour at different positions along PCA2 (Fig. 2b) highlights that colour variation along the pale–dark gradient co-occurs with systematic shifts in redness and saturation captured by PCA2.

### 3.2 Geographic structure of ant colour assemblages

Mean colour composition of ant assemblages exhibited strong geographic structure at the global scale (Fig. 3). Along the pale–dark axis (PCA1), assemblages were on average paler at higher latitudes, especially in North America and Europe, whereas darker assemblages predominated at low latitudes (Fig. 3a). Very dark mean PCA1 values were concentrated near the Equator, as reflected by a pronounced peak in darkness predicted by the generalized additive model (GAM; edf = 5.48, F = 27.04, P < 0.001; Fig. 3b), with latitude explaining 23.7% of the deviance in assemblage-level colour composition. In contrast, several subtropical and arid regions, including large parts of the Sahara and central to western Australia, were characterised by relatively pale assemblages.

**Figure 3.**
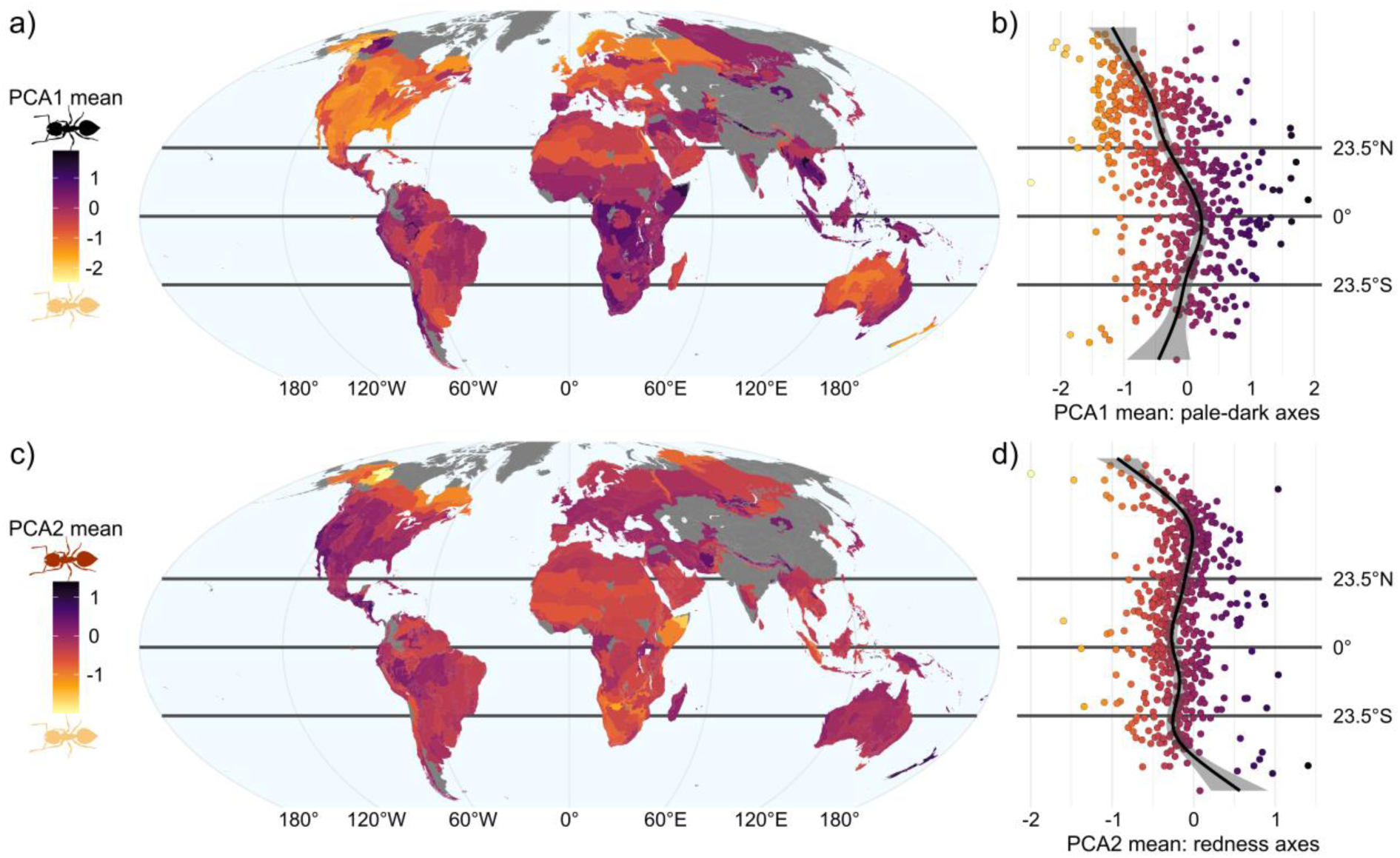
Geographic patterns of mean ant colour variation across ecoregions. (a) Global distribution of the mean PCA1 score (pale–dark colour axis) across terrestrial ecoregions. For each ecoregion, PCA1 values were averaged across all ant species present. Colours indicate the mean position of assemblages along PCA1, with lower values corresponding to paler assemblages and higher values to darker assemblages. (b) Latitudinal variation in mean PCA1 values across ecoregions. Points represent ecoregion centroids projected onto latitude and are coloured according to mean PCA1 values. The solid line shows the fitted relationship from a generalized additive model (GAM), with shaded areas indicating 95% confidence intervals. (c–d) Identical to (a–b), but for mean PCA2 values (redness axis). Horizontal lines indicate (North to South) the Tropic of Cancer, the Equator, and the Tropic of Capricorn, shown as reference boundaries of the tropical belt. Dark grey areas indicate ecoregions which we consider insufficiently sampled.

Geographic patterns along the redness axis (PCA2) were less spatially structured (Fig. 3c). Although mean PCA2 values varied significantly with latitude (GAM; edf = 7.54, F = 11.14, P < 0.001, Fig. 3d), the proportion of deviance explained was lower (14.5%). At the continental scale, assemblages were on average less red across much of Africa, whereas redder assemblages were more common in North and Central America, central Europe, and parts of Australia. Regionally low mean PCA2 values were observed at high northern latitudes in North America, contributing to the overall shape of the latitudinal trend.

### 3.3 Environmental and phylogenetic correlates of ant colour assemblages

We examined how the mean colour composition of ant assemblages along the pale–dark and redness axes covaried with climatic and environmental predictors, while accounting for phylogenetic and other biotic variables (Fig. 4). Spatial sdmTMB models accounting for residual spatial autocorrelation identified a limited set of robust correlates of assemblage colour composition (full model outputs in Tables S1 and S2).

**Figure 4.**
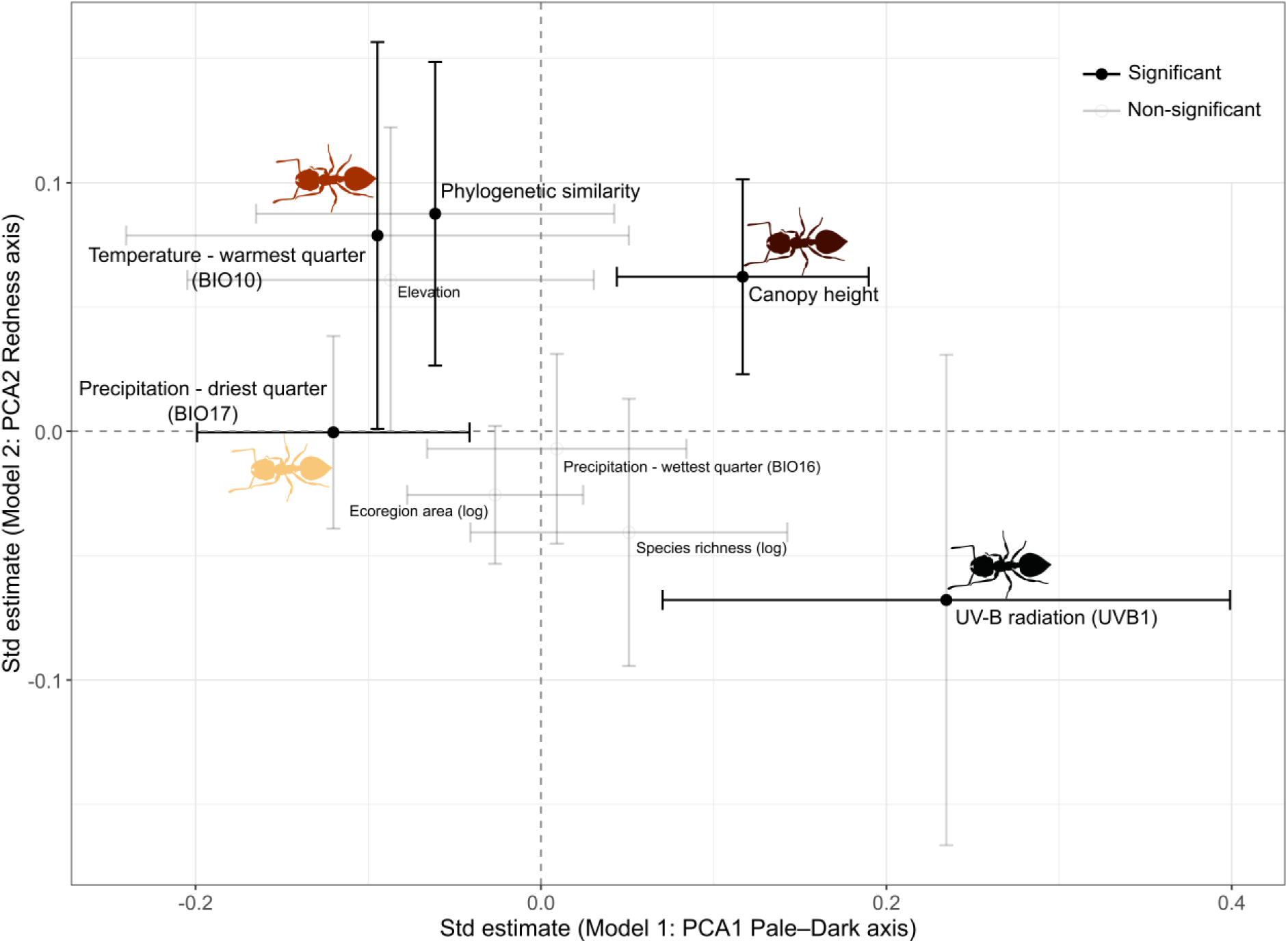
Environmental and phylogenetic correlates of ant colour assemblages. Bivariate forest plot showing standardised fixed-effect estimates from two spatial sdmTMB models relating ant assemblage mean colour composition to climatic, environmental and biotic predictors. The x-axis shows estimates from Model 1 predicting mean PCA1 values (pale–dark axis), and the y-axis shows estimates from Model 2 predicting mean PCA2 values (redness axis). Points represent coefficient estimates for the same predictors in both models, with horizontal (PCA1) and vertical (PCA2) error bars indicating 95% confidence intervals. Predictors were centred and scaled prior to analysis; estimates therefore represent the change in the response per one standard deviation increase in the predictor. Error bars are shown in black when confidence intervals exclude zero and in grey when they overlap zero. Dashed vertical and horizontal lines indicate zero effects.

Along the pale–dark axis (PCA1), darker assemblages (higher mean PCA1 values) were associated with higher UV-B radiation (standardised estimate = 0.235, 95% CI [0.070, 0.399]) and greater canopy height (0.117, [0.044, 0.190]). In contrast, mean PCA1 values decreased with increasing precipitation in the driest quarter (BIO17; −0.120, [−0.199, −0.041]), indicating paler assemblages in wetter dry-season environments. Other predictors, including temperature, species richness, phylogenetic similarity, ecoregion area and elevation, showed weak or non-significant associations with PCA1 (Fig. 4).

Redder assemblages (higher mean PCA2 values) were associated with greater phylogenetic similarity (0.088, [0.026, 0.149]), warmer temperatures in the warmest quarter (BIO10; 0.079, [0.001, 0.157]) and greater canopy height (0.062, [0.023, 0.101]). Evidence for effects of ecoregion area and elevation on PCA2 was comparatively weak, with confidence intervals close to zero, and the remaining predictors were not significantly associated with redness (Fig. 4).

Overall, fixed effects explained substantially more variation in the pale–dark axis than in the redness axis (R² = 0.193 vs 0.061; n = 586 assemblages). Residual spatial autocorrelation was low and non-significant for both models (Table S1-S2), indicating that spatial structure was adequately captured.

## 4 Discussion

Our global analysis shows that ant colouration at the assemblage level cannot be explained by a single biogeographical rule. Instead, it reflects the combined effects of multiple environmental drivers that act largely independently on the two main axes of colour variation: darkness and redness.

Because our analyses are conducted at the assemblage level, these patterns most likely reflect assemblage-level environmental filtering rather than evolutionary responses within species. In other words, environmental conditions may influence the mean colour composition of local ant assemblages through species turnover and lineage replacement, rather than through adaptive colour change within individual species. Nevertheless, identifying consistent relationships between environmental gradients and assemblage-level colour composition provides important insight into the ecological processes shaping large-scale colour patterns.

### 4.1 Darkness

Across ant assemblages at the global scale, we found no support for the prediction that darker assemblages should occur in warmer or more humid environments, as predicted by the complex Gloger’s rule (Fig. 1; Fig. 4). Instead, darker assemblages were consistently associated with lower precipitation during the driest quarter (BIO17), indicating increased melanisation in regions experiencing stronger seasonal water limitation (Fig. 4; Fig. S4; Table S1), consistent with the desiccation-resistance hypothesis. While this mechanism has been documented at local or regional scales (Davis & Moyle 2019; Parkash *et al*. 2008a, b, 2009b, a, 2012; Rajpurohit *et al*. 2008; Ramniwas & Kajla 2012), our study provides the first test at a global scale. These results contrast with global-scale patterns reported in passerine birds, where darker plumage is consistently associated with warm and humid climates (Delhey *et al*. 2019).

The classical association between darkness, warm climates and/or humidity has also often been interpreted indirectly through correlations with ultraviolet radiation, as warmer and more humid regions tend to coincide with higher UV exposure. By explicitly disentangling the effects of temperature, precipitation and UV-B radiation, we identified UV-B exposure as the main driver of assemblage-level darkness in ants. Consistent with the pronounced equatorial peak in darkness observed at the global scale (Fig. 3a, b), assemblages were consistently darker in regions with higher UV-B radiation (Fig. 4; Fig. S4; Table S1), consistent with the photoprotection hypothesis (Fig. 1) as a key mechanism underlying global patterns of melanisation in ants. Importantly, both photoprotection and desiccation resistance have previously been invoked to explain colour variation in ants at fine spatial scales. For example, along vertical vegetation strata, canopy-dwelling species tend to be darker than species inhabiting lower strata, which are typically more humid and less exposed to UV radiation (Law *et al*. 2020). Our results show that these mechanisms are not restricted to local ecological contexts but also scale up to explain global patterns of darkness across ant assemblages.

Apparent inconsistencies among previous studies may partly reflect differences in the level of biological organisation analysed. Previous work on ant colouration has focused almost exclusively on eumelanin-based darkness (Bishop *et al*. 2016; Bochenski *et al*. 2025; Idec *et al*. 2023; Klunk *et al*. 2022; Law *et al*. 2020; Skaldina *et al*. 2018; Skaldina & Sorvari 2017a, b), with only a few exceptions addressing other colour components in specific contexts such as mimicry (Schifani *et al*. 2024; Wagner *et al*. 2025; Wagner & Csősz 2025). A global interspecific analysis found no support for classical biogeographical predictions of dark colouration (Idec et al. 2023), and a separate global analysis focusing on the genus Pheidole similarly detected only weak colour–environment relationships (Klunk et al. 2022). In contrast, assemblage-level studies have detected significant colour-environment relationships along specific environmental gradients. For example, altitudinal gradient studies revealed darker assemblages at higher elevations, consistent with a combination of photoprotection and thermal melanism (Bishop *et al*. 2016), whereas studies along vertical vegetation strata detected signatures consistent with photoprotection and desiccation resistance (Law *et al*. 2020). Together, these contrasting results suggest that colour–environment relationships in ants may be more readily detected at the assemblage level, where environmental filtering acts on the mean colour composition of local communities. The weak effect of elevation in our global models likely reflects this scale dependency and the composite nature of elevation, which integrates multiple environmental drivers. While altitudinal gradient studies capture steep local environmental transitions over short distances, our analyses use mean ecoregion elevation and therefore reflect broader topographic context. Thermal melanism may thus operate primarily along local gradients, while being overshadowed at larger scales by stronger drivers such as UV radiation and moisture limitation.

Finally, assemblages occurring in regions with higher forest canopies were consistently darker (Fig. 4; Fig. S4; Table S1). Forest environments have often been invoked as an indirect explanation for the classical Gloger pattern linking darkness to warm and humid climates (Delhey 2019), but by explicitly separating climatic variables from vegetation structure, our results suggest that habitat itself may act as a direct driver of melanisation. This relationship is consistent with previously proposed mechanisms associated with forest environments, including enhanced camouflage in low-luminance and visually complex habitats (e.g., background matching (Merilaita *et al*. 2017)) and increased pathogen pressure (Fig. 1; Delhey 2019). It also echoes recent global findings in squamates highlighting habitat openness as a major determinant of colour variation (Goldenberg *et al*. 2025).

### 4.2 Redness

The complex version of Gloger’s rule predicts increased pheomelanin-based redness in warmer environments (Delhey 2019; Fig. 1), a prediction supported by our global analyses (Fig. 4; Fig. S4; Table S2), with assemblages occurring in regions with higher temperatures during the warmest quarter being consistently redder. Although the complex version of Gloger’s rule also predicts increased redness under dry conditions, we did not detect an effect of precipitation variables on redness (Fig. 4), indicating that temperature may represent the dominant climatic driver of redness in ants.

Habitat structure also contributed to variation in redness. Assemblages occurring in regions with taller forest canopies were consistently redder, indicating that vegetation structure may influence the two main axes of colour variation simultaneously (Fig. 4; Fig. S4). This pattern is consistent with sensory-environment constraints imposed by forest light conditions, under which reddish-brown colouration may enhance crypsis or background matching (Endler 1993; Gomez & Thery 2007; Gomez & Théry 2004).

Notably, we found no evidence that ultraviolet radiation constrained redness (Fig. 4), despite predictions from the phototoxicity hypothesis that pheomelanin may incur oxidative costs under high UV exposure (Ito *et al*. 2018). This absence of effect suggests that potential phototoxic costs may be outweighed by other selective pressures, or that pheomelanin levels in ants remain within physiological ranges that limit phototoxic damage, at least when considering assemblage-level mean colouration. More generally, interpreting redness patterns in ants requires some caution because the biochemical basis of reddish colouration in this group remains poorly understood. Our conceptual framework is grounded in hypotheses developed for pheomelanin-based pigmentation (Fig. 1), yet it is not currently known whether pheomelanin is the primary pigment responsible for red hues in ants. Nevertheless, studies in other hymenopterans have shown that shifts between black and red colouration can arise from changes in the relative contribution of eumelanin and pheomelanin pigments (Hines *et al*. 2017), suggesting that similar mechanisms could plausibly operate in ants. As multiple pigments could contribute to red colouration in insects (Badejo *et al*. 2020), variation unrelated to pheomelanin-based mechanisms could introduce additional noise into macroecological patterns, potentially contributing to the lower explanatory power of the redness models compared with those for darkness (Tables S1–S2). In contrast, eumelanin is well established as the dominant pigment underlying dark colour variation across insects (Badejo *et al*. 2020) and is therefore very likely to represent the primary pigment involved in ants. Interestingly, the positive association between redness and temperature remains consistent with predictions derived from the complex version of Gloger’s rule despite this biochemical uncertainty.

Finally, phylogenetic similarity among assemblages also structured variation in redness (Fig. 4; Fig. S4, Table S2). Assemblages with more similar phylogenetic composition (Fig. S5) tended to exhibit similar levels of redness, suggesting that evolutionary history and lineage composition influence large-scale colour patterns alongside climatic and environmental drivers. This effect likely reflects differences in pigment expression among major ant clades, combined with the geographic structuring of evolutionary lineages across regions.

### 4.3 Beyond Gloger’s rule

Overall, our results indicate that large-scale colour variation in ants does not conform to expectations derived from the complex version of Gloger’s rule. Instead, assemblage-level colour composition is primarily shaped by alternative mechanisms, notably photoprotection, desiccation resistance, and habitat-related camouflage. Only redness showed partial agreement with the complex framework through its association with temperature, although this effect remained comparatively weak and explained substantially less variation than patterns observed for darkness (Tables S1–S2). An important insight emerging from our analyses is that the two main axes of ant colour variation respond differently to environmental and evolutionary processes. Darkness was primarily associated with contemporary environmental drivers, particularly ultraviolet radiation and seasonal water limitation, whereas redness showed a stronger signature of phylogenetic structure among assemblages. This asymmetry suggests that different pigmentary components may be subject to distinct constraints, with eumelanin-based darkness appearing more environmentally labile, while reddish colouration may be more strongly shaped by evolutionary history and lineage-specific pigmentation pathways. A similar asymmetry has been reported in lagomorphs, where reddish pigmentation shows stronger phylogenetic structuring than darker pigmentation (Cerezer *et al*. 2025).

This asymmetry highlights that large-scale colour patterns cannot always be explained by a single ecogeographical rule, but may instead emerge from the interaction between environmental filtering and the evolutionary history of pigment expression across lineages. Empirical support for the complex formulation of Gloger’s rule is already heterogeneous across vertebrate clades (Fig. 5), with different taxa supporting different components of the framework. The even weaker correspondence observed in ants suggests a stronger departure from Gloger’s predictions than the variation reported among vertebrate groups themselves, indicating that mechanisms inferred from vertebrate systems may not generalise across taxa with contrasting physiology and ecological constraints. More broadly, our findings highlight that large-scale colour variation can result from multiple selective pressures acting on distinct pigmentary axes. Some drivers act independently (e.g. ultraviolet radiation affecting darkness), whereas others act simultaneously on both axes (e.g. vegetation structure influencing both darkness and redness). Because colour directly mediates interactions between organisms and their environment (Cuthill *et al*. 2017), our study highlights how multiple ecological constraints jointly shape large-scale colour variation in a critical insect group (Schultheiss *et al*. 2022).

**Figure 5.**
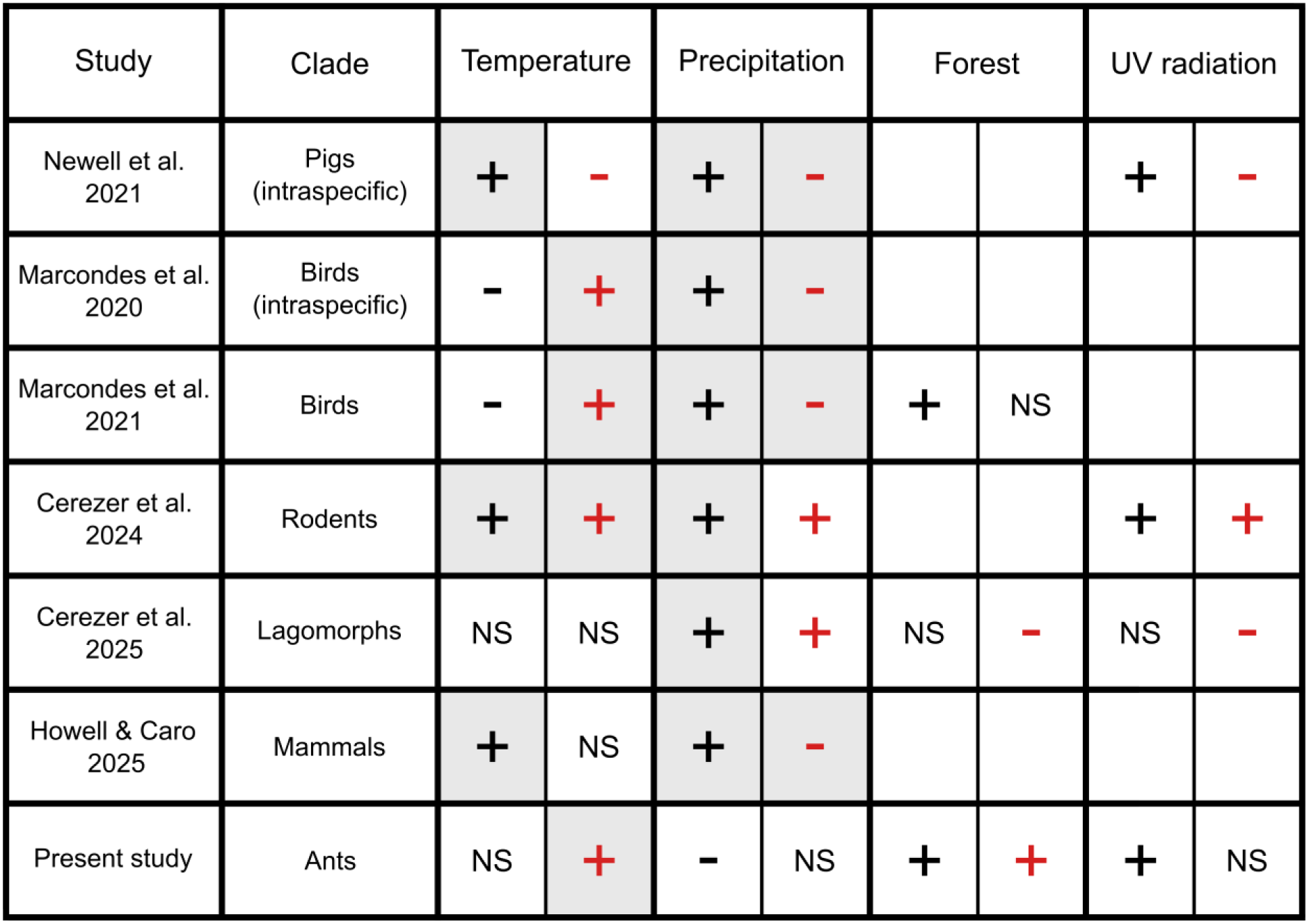
Summary of studies that analysed variation in both dark (eumelanin-based) and red (pheomelanin-based) colour components within the same analytical framework. Columns indicate the main environmental predictors. Black symbols refer to eumelanin-based darkness and red symbols to pheomelanin-based redness. Plus (+) and minus (−) symbols indicate positive and negative relationships. Grey cells indicate relationships in the direction predicted by the complex formulation of Gloger’s rule (Fig. 1). NS indicates non-significant relationships, and empty cells indicate predictors that were not tested in the corresponding study.

### 4.4 Limitations and future directions

Although our conceptual framework relies on melanin-based pigmentation (eumelanin and pheomelanin), the biochemical basis of colour variation in ants remains unknown. The detection of consistent macroecological patterns nevertheless suggests that either melanin is involved, at least partially, or alternative pigments respond to similar environmental constraints. Future work combining chemical analyses of cuticular pigments with quantitative colour measurements, as well as experimental studies linking UV and desiccation resistance to melanisation across species, as done in *Drosophila* (Lotfy *et al*. 2024; Rajpurohit *et al*. 2008; Ramniwas *et al*. 2012), would help clarify the mechanisms underlying darkness and redness variation.

Because colour measurements were derived from photographs rather than reflectance spectrometry, variation in imaging conditions could introduce noise. However, colour estimates from heterogeneous photographic datasets can closely match spectrometry when sample sizes are sufficient (12–14 images per taxon; Laitly *et al*. 2021). Although our per-species sample sizes were often lower, AntWeb images follow a highly standardized acquisition protocol (Idec *et al*. 2023), and manual selection of representative pixels while avoiding overexposed or shadowed regions further reduces illumination artefacts. Indeed, human observers can partially discount illumination changes and specular reflections when estimating surface appearance, through colour constancy and related perceptual mechanisms (Nohira & Nagai 2023; Schultz *et al*. 2006; Toscani *et al*. 2017). Although the AntWeb imaging pipeline follows a standardized protocol (Idec *et al*. 2023), photographs are taken with different camera systems by different users, which may introduce additional noise. However, such variation is unlikely to be structured and should therefore act as random noise, increasing the risk of Type II errors rather than generating the systematic patterns revealed here.

Finally, because our measurements are based on standard photography, they capture only variation within the visible spectrum. Extending analyses to other wavelengths would provide important insights: ultraviolet reflectance may show even stronger associations with UV-B compare to darkness, whereas near-infrared reflectance, which represents more than half of the solar radiant energy reaching the Earth’s surface, should primarily reflect thermoregulatory properties (Stuart-Fox *et al*. 2017).

### 4.5 Conclusions

Our results reveal that the two main axes of ant colour variation respond to different processes: darkness closely tracks environmental gradients, whereas redness retains a stronger phylogenetic signature. This asymmetry suggests that pigmentary traits may differ in their evolutionary lability, highlighting the need to consider multiple dimensions of colour when investigating large-scale ecological patterns.

## Acknowledgements

We gratefully acknowledge all those who contributed to the collection, imaging of the specimens and occurrence records used in this study, including the many contributors who helped build and expand AntWeb. We also thank Alexandros Topaloudis for his help with data curation. We further acknowledge the financial support of the Canton of Vaud, Switzerland.

## Statement of authorship

S.O., C.B., and T.K. conceived the study. T.K., S.O., C.B., and E.P. developed the methodology. T.K. performed the analyses and produced the figures. B.L.F. provided data resources, and S.O. curated the data. T.K. wrote the first draft of the manuscript, and all authors (T.K., B.L.F., E.P., C.B., and S.O.) contributed substantially to revisions. The project was supervised equally by S.O. and C.B.

## Data accessibility statement

Data and code supporting the results of this study will be archived in the Dryad Digital Repository upon acceptance.

---

**Figure S1.**
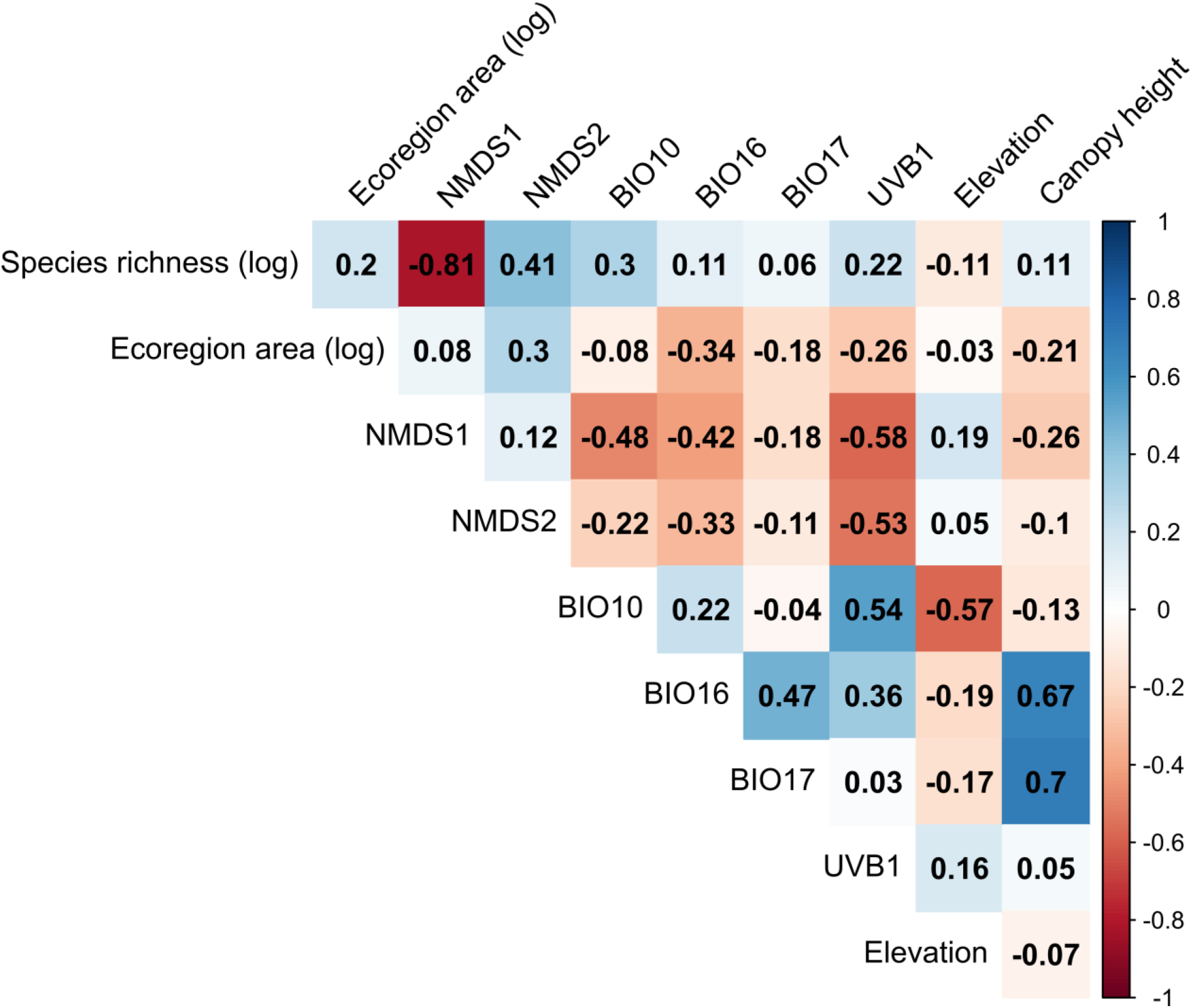
Correlation structure among explanatory variables. Pairwise Pearson correlation coefficients among the ten predictors initially assembled for the spatial models. All variables were mean-centred and scaled to unit variance prior to analysis. NMDS1 and NMDS2 correspond to the first two axes of a non-metric multidimensional scaling (NMDS) ordination based on pairwise phylogenetic distances among species assemblages. Climatic variables include temperature of the warmest quarter (BIO10), precipitation of the wettest quarter (BIO16) and precipitation of the driest quarter (BIO17), extracted from WorldClim v2.1. Elevation represents mean ecoregion elevation. UV-B radiation (UVB1) corresponds to mean annual UV-B radiation. Species richness represents the number of ant species per assemblage (ecoregion), log-transformed prior to scaling. Ecoregion area corresponds to the total area of each ecoregion (log-transformed). Colours indicate the strength and direction of correlations (blue = positive, red = negative).

**Figure S2.**
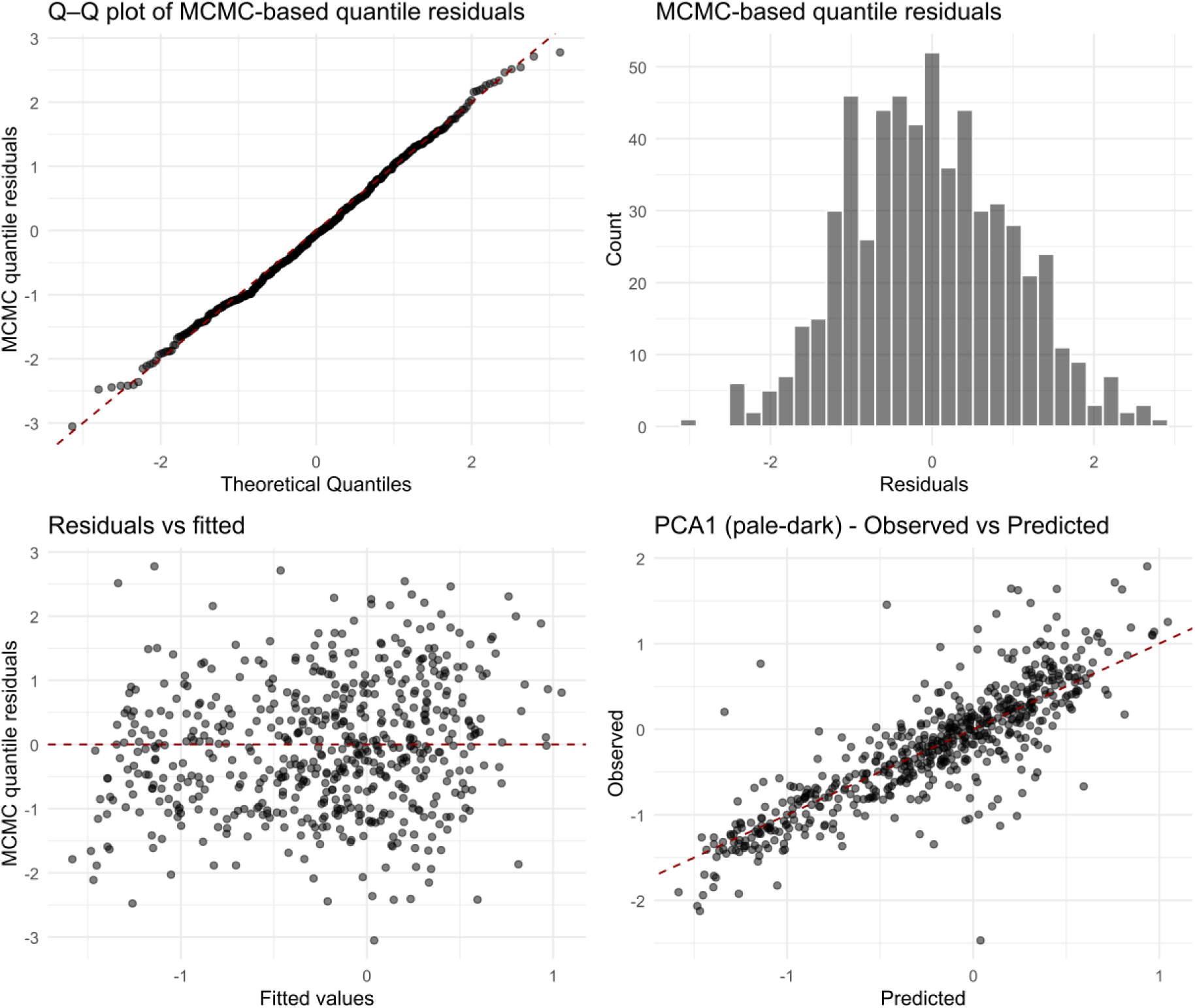
Diagnostic plots for the spatial model of the pale–dark colour axis (PCA1). Diagnostic plots for the spatial sdmTMB model fitted to mean PCA1 values using a Student-t error distribution. The upper-left panel shows a Q–Q plot of MCMC-based quantile residuals against theoretical normal quantiles, with the dashed line indicating the 1:1 relationship. The upper-right panel displays the distribution of MCMC-based quantile residuals. The lower-left panel shows quantile residuals plotted against fitted values, and the lower-right panel presents observed versus predicted PCA1 values, with the dashed line indicating the 1:1 relationship. Overall, residuals show no strong deviations from model assumptions, indicating an adequate fit of the spatial model.

**Figure S3.**
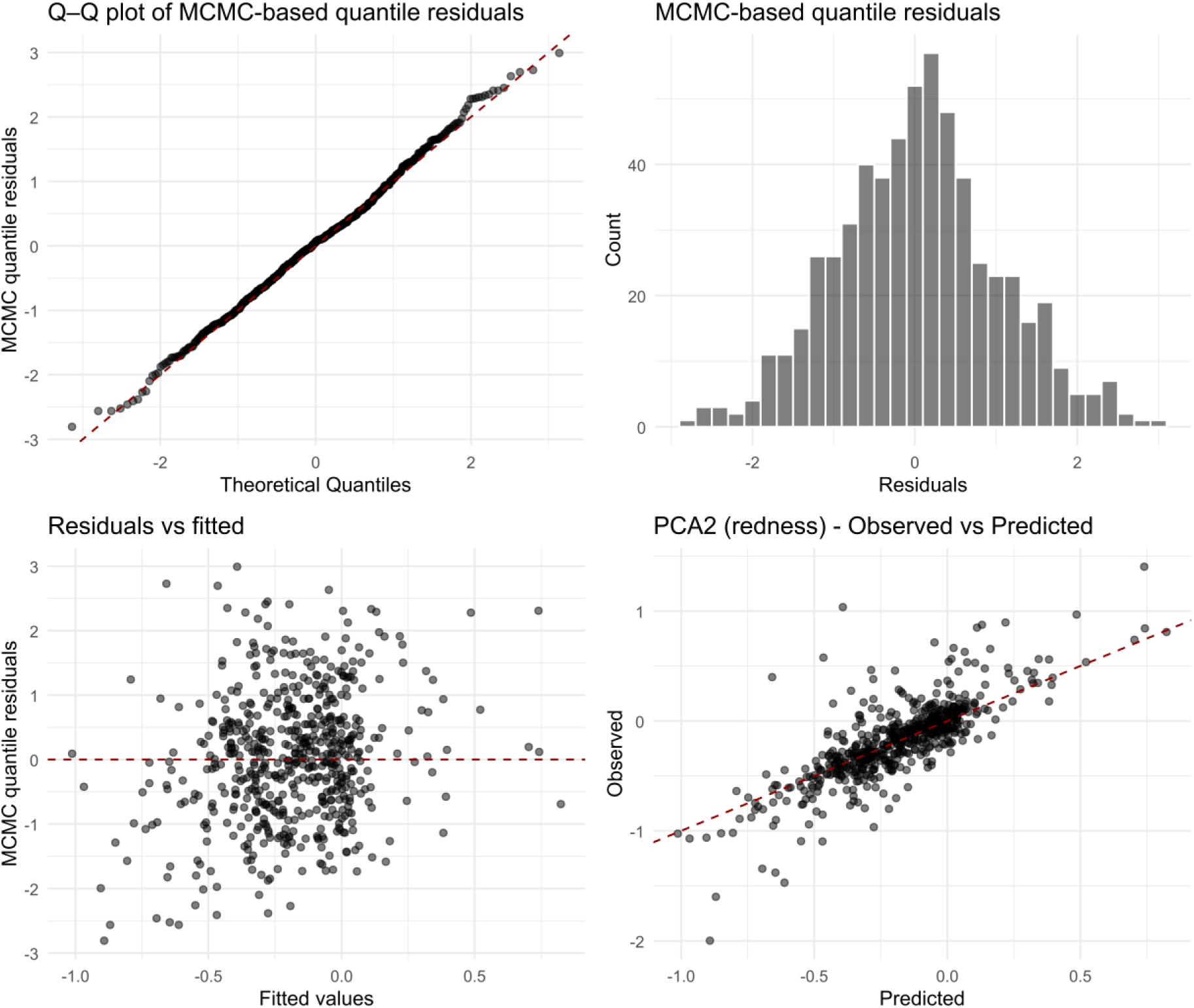
Diagnostic plots for the spatial model of the redness axis (PCA2). Diagnostic plots for the spatial sdmTMB model fitted to mean PCA2 values using a Student-t error distribution. The upper-left panel shows a Q–Q plot of MCMC-based quantile residuals against theoretical normal quantiles, with the dashed line indicating the 1:1 relationship. The upper-right panel displays the distribution of MCMC-based quantile residuals. The lower-left panel shows quantile residuals plotted against fitted values, and the lower-right panel presents observed versus predicted PCA2 values, with the dashed line indicating the 1:1 relationship. Overall, residuals show no strong deviations from model assumptions, indicating an adequate fit of the spatial model.

**Figure S4:**
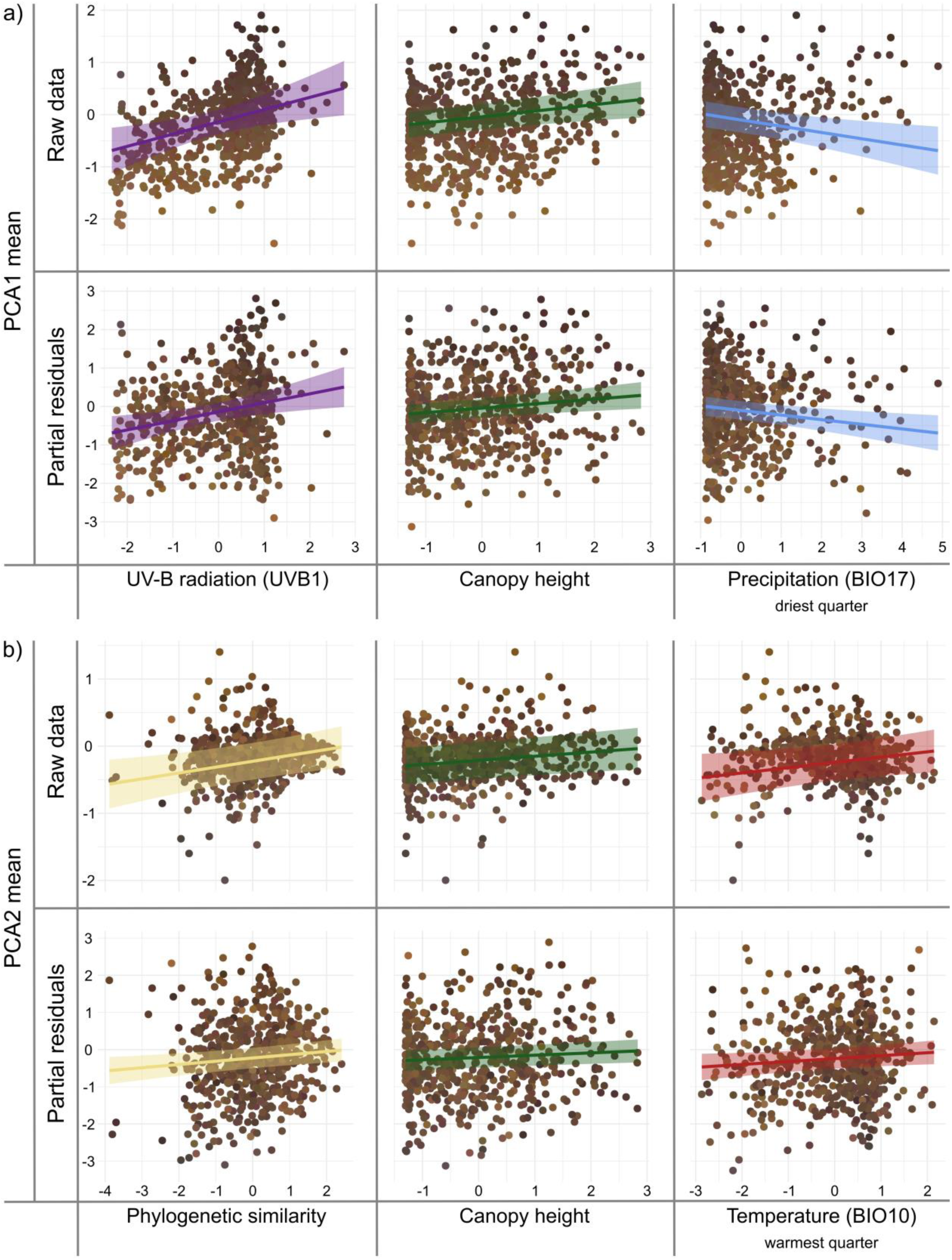
Conditional effects from spatial sdmTMB models for (a) PCA1 (pale-dark axis) and (b) PCA2 (redness axis), shown only for predictors whose 95% confidence interval (or a more stringent confidence interval) does not include zero. For each predictor, the same fitted conditional relationship (solid line) and its confidence band are displayed in both rows: the top row overlays this relationship on the raw PCA scores, whereas the bottom row overlays it on the corresponding partial residuals. Each point represents the mean PCA1 or PCA2 value calculated for one of the 586 ant assemblages worldwide and is coloured using the average thoracic colour of a real ant species. For each assemblage, its mean PCA1–PCA2 coordinates are projected into the ant colour PCA space (derived from ∼10,400 species), and the colour of the nearest species in that space is assigned.

**Figure S5.**
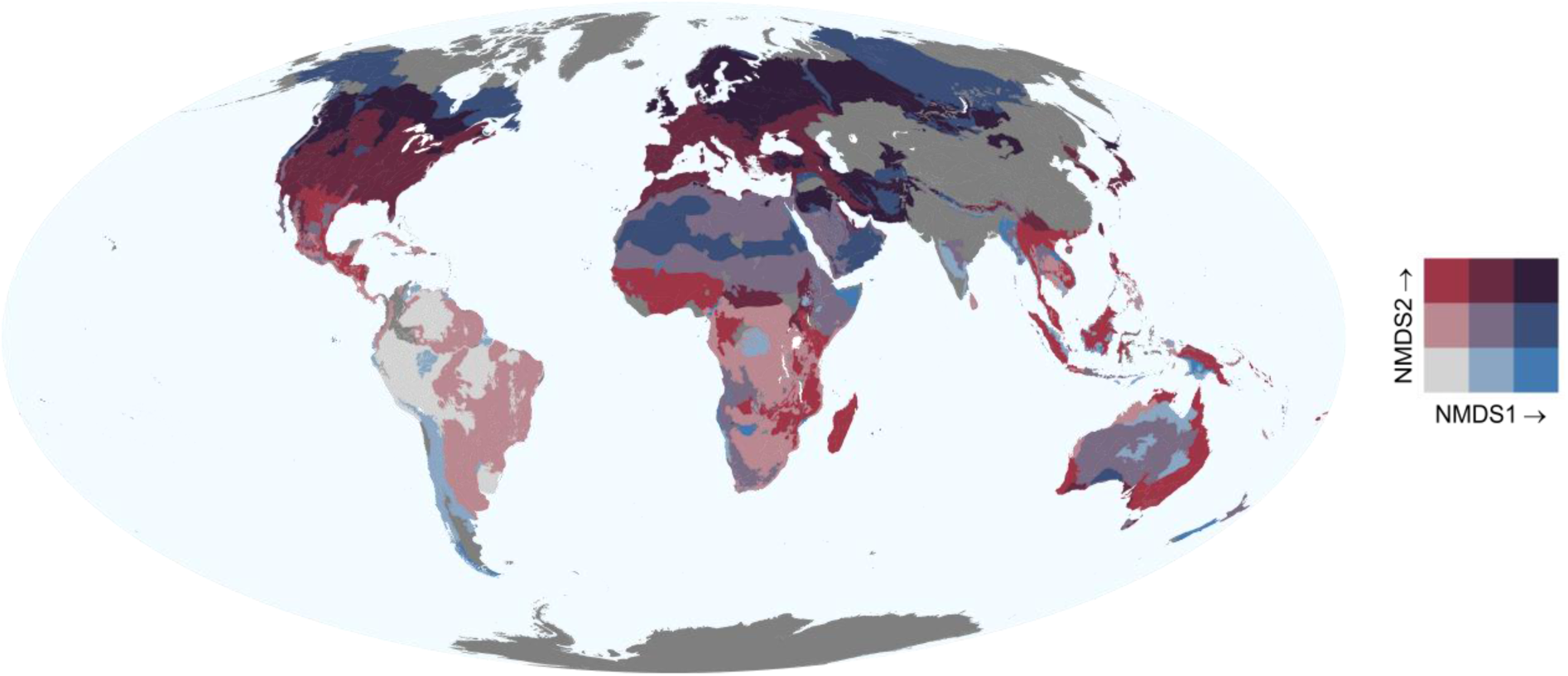
Global distribution of phylogenetic similarity among ant assemblages. Bivariate map showing the spatial distribution of the first two axes of the non-metric multidimensional scaling (NMDS) ordination derived from the phylogenetic similarity matrix among ecoregions. Colours represent the joint position of each ecoregion along NMDS1 (horizontal gradient) and NMDS2 (vertical gradient), such that ecoregions with similar colours have more similar phylogenetic composition. NMDS2 was retained in the main analyses as an index of phylogenetic similarity among assemblages because NMDS1 was strongly correlated with species richness and therefore excluded. Dark grey areas indicate ecoregions for which we have insufficient data.

**Table S1.** Fixed-effect estimates from the spatial SDM model explaining variation in PCA1 (Pale–Dark axis) using sdmTMB. Estimates correspond to regression coefficients and are reported with their standard errors (SE) and 90%, 95%, 99% and 99.9% confidence intervals (CI). Predictors were standardised prior to analysis and spatial structure was modelled using a mesh-based random field. Stars indicate the most stringent confidence interval that does not include zero (* = 90% CI, ** = 95% CI, *** = 99% CI, **** = 99.9% CI); no star indicates that all confidence intervals include zero.

| PCA1: Pale-Dark |  |  |  |  |  |  |  |
| --- | --- | --- | --- | --- | --- | --- | --- |
| Predictor | Estimate | SE | CI 90% | CI 95% | CI 99% | CI 99.9% | Sig |
| (Intercept) | -0.177 | 0.117 | [-0.369, 0.015] | [-0.406, 0.052] | [-0.478, 0.124] | [-0.562, 0.208] |  |
| Species richness (log) | 0.051 | 0.047 | [-0.026, 0.128] | [-0.041, 0.143] | [-0.070, 0.171] | [-0.103, 0.205] |  |
| Phylogenetic similarity | -0.061 | 0.053 | [-0.148, 0.026] | [-0.165, 0.042] | [-0.198, 0.075] | [-0.235, 0.113] |  |
| Ecoregion area (log) | -0.027 | 0.026 | [-0.069, 0.016] | [-0.077, 0.024] | [-0.093, 0.040] | [-0.112, 0.059] |  |
| Temperature - warmest quarter (BIO10) | -0.095 | 0.074 | [-0.217, 0.027] | [-0.240, 0.051] | [-0.286, 0.097] | [-0.339, 0.150] |  |
| Precipitation - wettest quarter (BIO16) | 0.009 | 0.038 | [-0.054, 0.072] | [-0.066, 0.084] | [-0.090, 0.108] | [-0.117, 0.135] |  |
| **Precipitation - driest quarter (BIO17)** | **-0.120** | 0.040 | [-0.186, -0.054] | [-0.199, -0.041] | **[-0.224, -0.017]** | [-0.253, 0.012] | **\*\*** |
| **UV-B radiation (UVB1)** | **0.235** | 0.084 | [0.097, 0.373] | [0.070, 0.399] | **[0.019, 0.451]** | [-0.041, 0.511] | **\*\*** |
| Elevation | -0.087 | 0.060 | [-0.186, 0.012] | [-0.205, 0.031] | [-0.242, 0.067] | [-0.285, 0.110] |  |
| **Canopy height** | **0.117** | 0.037 | [0.056, 0.178] | [0.044, 0.190] | **[0.021, 0.213]** | [-0.006, 0.239] | **\*\*** |
| R² (fixed effects) = 0.193; n = 586. |  |  |  |  |  |  |  |
| Moran's I = -0.0035 (p-value = 0.55) |  |  |  |  |  |  |  |

**Table S2.** Fixed-effect estimates from the spatial SDM model explaining variation in PCA2 (redness axis) using sdmTMB. Estimates correspond to regression coefficients and are reported with their standard errors (SE) and 90%, 95%, 99% and 99.9% confidence intervals (CI). Predictors were standardised prior to analysis and spatial structure was modelled using a mesh-based random field. Stars indicate the most stringent confidence interval that does not include zero (* = 90% CI, ** = 95% CI, *** = 99% CI, **** = 99.9% CI); no star indicates that all confidence intervals include zero.

| PCA2: Redness |  |  |  |  |  |  |  |
| --- | --- | --- | --- | --- | --- | --- | --- |
| Predictor | Estimate | SE | CI 90% | CI 95% | CI 99% | CI 99.9% | Sig |
| (Intercept) | -0.186 | 0.136 | [-0.410, 0.039] | [-0.453, 0.082] | [-0.537, 0.166] | [-0.635, 0.263] |  |
| Species richness (log) | -0.041 | 0.027 | [-0.086, 0.004] | [-0.094, 0.013] | [-0.111, 0.030] | [-0.131, 0.050] |  |
| **Phylogenetic similarity** | **0.088** | 0.031 | [0.036, 0.139] | [0.026, 0.149] | **[0.007, 0.168]** | [-0.015, 0.190] | **\*\*** |
| Ecoregion area (log) | -0.026 | 0.014 | [-0.049, -0.002] | [-0.053, 0.002] | [-0.062, 0.011] | [-0.072, 0.021] | . |
| **Temperature - warmest quarter (BIO10)** | **0.079** | 0.040 | [0.013, 0.144] | **[0.001, 0.157]** | [-0.023, 0.181] | [-0.052, 0.209] | **\*** |
| Precipitation - wettest quarter (BIO16) | -0.007 | 0.019 | [-0.039, 0.025] | [-0.045, 0.031] | [-0.057, 0.043] | [-0.071, 0.057] |  |
| Precipitation - driest quarter (BIO17) | 0.000 | 0.020 | [-0.033, 0.032] | [-0.039, 0.038] | [-0.051, 0.051] | [-0.065, 0.065] |  |
| UV-B radiation (UVB1) | -0.068 | 0.050 | [-0.151, 0.015] | [-0.166, 0.031] | [-0.197, 0.062] | [-0.233, 0.098] |  |
| Elevation | 0.061 | 0.031 | [0.009, 0.112] | [-0.000, 0.122] | [-0.020, 0.142] | [-0.042, 0.164] | . |
| **Canopy height** | **0.062** | 0.020 | [0.029, 0.095] | [0.023, 0.101] | **[0.011, 0.114]** | [-0.004, 0.128] | **\*\*** |
| R<sup>2</sup> (fixed effects) = 0.061; n = 586. |  |  |  |  |  |  |  |
| Moran's I = -0.0059 (p-value = 0.61) |  |  |  |  |  |  |  |

## References

1. Andersen, S.O. (2012). Cuticular Sclerotization and Tanning. In: Insect Molecular Biology and Biochemistry. Elsevier, pp. 167–192.

2. Anderson, S.C., Ward, E.J., English, P.A., Barnett, L.A.K. & Thorson, J.T. (2025). sdmTMB: An R Package for Fast, Flexible, and User-Friendly Generalized Linear Mixed Effects Models with Spatial and Spatiotemporal Random Fields. Journal of Statistical Software, 115, 1–46.

3. Arnan, X., Cerdá, X. & Retana, J. (2015). Partitioning the impact of environment and spatial structure on alpha and beta components of taxonomic, functional, and phylogenetic diversity in European ants. PeerJ, 3, e1241.

4. Badejo, O., Skaldina, O., Gilev, A. & Sorvari, J. (2020). Benefits of insect colours: a review from social insect studies. Oecologia, 194, 27–40.

5. Beckmann, M., Václavík, T., Manceur, A.M., Šprtová, L., Wehrden, H. von, Welk, E., et al. (2014). glUV: A global UV-B radiation data set for macroecological studies. Methods in Ecology and Evolution, 5, 372–383.

6. Bishop, T.R., Robertson, M.P., Gibb, H., Rensburg, B.J. van, Braschler, B., Chown, S.L., et al. (2016). Ant assemblages have darker and larger members in cold environments. Global Ecology and Biogeography, 25, 1489–1499.

7. Bochenski, W.S., Vicente, R.E., Bishop, T.R. & Izzo, T.J. (2025). Ant brightness and size influence the seasonal vegetation use by ground-dwelling ants (Hymenoptera: Formicidae) in an Amazonia-Cerrado transition zone. Ecological Entomology, 50, 1082–1095.

8. Bogert, C.M. (1949). Thermoregulation in Reptiles, A Factor in Evolution. Evolution, 3, 195–211.

9. Cavender-Bares, J., Kozak, K.H., Fine, P.V.A. & Kembel, S.W. (2009). The merging of community ecology and phylogenetic biology. Ecology Letters, 12, 693–715.

10. Cerezer, F.O., Campos, A.B. & Cáceres, N.C. (2025). Ecological and Evolutionary Predictors of Dorsal Colouration in Lagomorphs. Journal of Biogeography, 52, e70016.

11. Cerezer, F.O., Campos, A.B., Dambros, C.S., Maestri, R., Bubadué, J.M. & Cáceres, N.C. (2024). Rodents show darker and redder coloration in warm and rainy environments. Global Ecol Biogeogr, 33, 426–438.

12. Clusella Trullas, S., van Wyk, J.H. & Spotila, J.R. (2007). Thermal melanism in ectotherms. Journal of Thermal Biology, 32, 235–245.

13. Clusella-Trullas, S. & Nielsen, M. (2020). The evolution of insect body coloration under changing climates. Current Opinion in Insect Science. 41, 25–32.

14. Cooney, C.R., He, Y., Varley, Z.K., Nouri, L.O., Moody, C.J.A., Jardine, M.D., et al. (2022). Latitudinal gradients in avian colourfulness. Nature Ecology and Evolution, 6, 622–629.

15. Cuthill, I.C., Allen, W.L., Arbuckle, K., Caspers, B., Chaplin, G., Hauber, M.E., et al. (2017). The biology of color. Science, 357, eaan0221.

16. Davis, J.S. & Moyle, L.C. (2019). Desiccation resistance and pigmentation variation reflects bioclimatic differences in the Drosophila americana species complex. BMC Evol Biol, 19, 204.

17. Delhey, K. (2017). Gloger’s rule. Current Biology, 27, R689–R691.

18. Delhey, K. (2019). A review of Gloger’s rule, an ecogeographical rule of colour: definitions, interpretations and evidence. Biological Reviews, 94, 1294–1316.

19. Delhey, K., Dale, J., Valcu, M. & Kempenaers, B. (2019). Reconciling ecogeographical rules: rainfall and temperature predict global colour variation in the largest bird radiation. Ecology Letters, 22, 726–736.

20. Dinerstein, E., Olson, D., Joshi, A., Vynne, C., Burgess, N.D., Wikramanayake, E., et al. (2017). An Ecoregion-Based Approach to Protecting Half the Terrestrial Realm. BioScience, 67, 534–545.

21. Doré, M., Borowiec, M.L., Branstetter, M.G., Camacho, G.P., Fisher, B.L., Longino, J.T., et al. (2025). Evolutionary history of ponerine ants highlights how the timing of dispersal events shapes modern biodiversity. Nat Commun, 16, 8297.

22. Dormann, C.F., Elith, J., Bacher, S., Buchmann, C., Carl, G., Carré, G., et al. (2013). Collinearity: a review of methods to deal with it and a simulation study evaluating their performance. Ecography, 36, 27–46.

23. Dray, S. & Dufour, A.-B. (2007). The ade4 package: implementing the duality diagram for ecologists. Journal of Statistical Software, 22, 1–20.

24. Dubovskiy, I.M., Whitten, M.M.A., Kryukov, V.Y., Yaroslavtseva, O.N., Grizanova, E.V., Greig, C., et al. (2013). More than a colour change: insect melanism, disease resistance and fecundity. Proc Biol Sci, 280, 20130584.

25. Ducrest, A.-L., Keller, L. & Roulin, A. (2008). Pleiotropy in the melanocortin system, coloration and behavioural syndromes. Trends in Ecology & Evolution, 23, 502–510.

26. Economo, E.P., Narula, N., Friedman, N.R., Weiser, M.D. & Guénard, B. (2018). Macroecology and macroevolution of the latitudinal diversity gradient in ants. Nat Commun, 9, 1778.

27. Endler, J.A. (1993). The Color of Light in Forests and Its Implications. Ecological Monographs, 63, 1–27.

28. Fick, S.E. & Hijmans, R.J. (2017). WorldClim 2: new 1-km spatial resolution climate surfaces for global land areas. International Journal of Climatology, 37, 4302–4315.

29. Fraenkel, G. & Rudall, K.M. (1940). A study of the physical and chemical properties of the insect cuticle. Proc. R. Soc. Lond. B, 129, 1–35.

30. Goldenberg, J., Bisschop, K., Lambert, J.W., Nicolaï, M.P.J., Etienne, R.S., D’Alba, L., et al. (2025). Habitat openness and squamate color evolution over deep time. Nat Commun, 16, 2625.

31. Gomez, D. & Théry, M. (2004). Influence of ambient light on the evolution of colour signals: comparative analysis of a Neotropical rainforest bird community. Ecology Letters, 7, 279–284.

32. Gomez, D. & Thery, M. (2007). Simultaneous Crypsis and Conspicuousness in Color Patterns: Comparative Analysis of a Neotropical Rainforest Bird Community. the american naturalist, 169, S42–S61.

33. Hines, H.M., Witkowski, P., Wilson, J.S. & Wakamatsu, K. (2017). Melanic variation underlies aposematic color variation in two hymenopteran mimicry systems. PLoS ONE, 12.

34. Hölldobler, B. & Wilson, E. (1990). The Ants. Harvard University Press, Cambridge, MA.

35. Howell, N. & Caro, T. (2025). Gloger’s Rule or Historical Conjecture? Tests in Mammals. Ecology and Evolution, 15, e71855.

36. Idec, J.H., Bishop, T.R. & Fisher, B.L. (2023). Using computer vision to understand the global biogeography of ant color. Ecography, 2023, e06279.

37. Ito, S., Wakamatsu, K. & Sarna, T. (2018). Photodegradation of eumelanin and pheomelanin and its pathophysiological implications. Photochem. Photobiol., 94, 409–420.

38. Kass, J.M., Guénard, B., Dudley, K.L., Jenkins, C.N., Azuma, F., Fisher, B.L., et al. (2022). The global distribution of known and undiscovered ant biodiversity. Sci. Adv., 8, eabp9908.

39. Klunk, C.L., Fratoni, R.O., Rivadeneira, C.D., Schaedler, L.M. & Perez, D.M. (2022). Climate and body size have differential roles on melanism evolution across workers in a worldwide ant genus. Oecologia, 199, 579–587.

40. Laitly, A., Callaghan, C.T., Delhey, K. & Cornwell, W.K. (2021). Is color data from citizen science photographs reliable for biodiversity research? Ecology and Evolution, 11, 4071–4083.

41. Lang, N., Jetz, W., Schindler, K. & Wegner, J.D. (2023). A high-resolution canopy height model of the Earth. Nature Ecology and Evolution, 7, 1778–1789.

42. Law, S.J., Bishop, T.R., Eggleton, P., Griffiths, H., Ashton, L. & Parr, C. (2020). Darker ants dominate the canopy: Testing macroecological hypotheses for patterns in colour along a microclimatic gradient. Journal of Animal Ecology, 89, 347–359.

43. Lindgren, F., Rue, H. & Lindström, J. (2011). An Explicit Link between Gaussian Fields and Gaussian Markov Random Fields: The Stochastic Partial Differential Equation Approach. J. R. Stat. Soc. Ser. B. Stat. Methodol., 73, 423–498.

44. Lotfy, M., Khattab, A., Shata, M., Alhasbani, A., Almesmari, A., Alsaeedi, S., et al. (2024). Destructive effects of UVC radiation on Drosophila melanogaster: Mortality, fertility, mutations, and molecular mechanisms. PLOS ONE, 19, e0303115.

45. Marcondes, R.S., Nations, J.A., Seeholzer, G.F. & Brumfield, R.T. (2021). Rethinking Gloger’s Rule: Climate, Light Environments, and Color in a Large Family of Tropical Birds (Furnariidae). The American Naturalist, 197, 592–606.

46. Marcondes, R.S., Stryjewski, K.F. & Brumfield, R.T. (2020). Testing the simple and complex versions of Gloger’s rule in the Variable Antshrike (Thamnophilus caerulescens, Thamnophilidae). Auk, 137, ukaa026.

47. Matte, A., Billen, J., Shit, P., Heinze, J. & Bernadou, A. (2025). Delayed maturation of the exoskeleton and muscle fibres in the ant Platythyrea punctata. Biological Journal of the Linnean Society, 144, blae025.

48. Merilaita, S., Scott-Samuel, N.E. & Cuthill, I.C. (2017). How camouflage works. Philos Trans R Soc Lond B Biol Sci, 372, 20160341.

49. Newell, C., Walker, H. & Caro, T. (2021). Pig pigmentation: testing Gloger’s rule. Journal of Mammalogy, 102, 1525–1535.

50. Nohira, H. & Nagai, T. (2023). Texture statistics involved in specular highlight exclusion for object lightness perception. Journal of Vision, 23, 1.

51. Oksanen, J., Simpson, G.L., Blanchet, F.G., Kindt, R., Legendre, P., Minchin, P.R., et al. (2024). vegan: Community Ecology Package.

52. Parkash, R., Chahal, J., Sharma, V. & Dev, K. (2012). Adaptive associations between total body color dimorphism and climatic stress-related traits in a stenothermal circumtropical Drosophila species. Insect Science, 19, 247–262.

53. Parkash, R., Rajpurohit, S. & Ramniwas, S. (2008a). Changes in body melanisation and desiccation resistance in highland vs. lowland populations of *D. melanogaster*. Journal of Insect Physiology, 54, 1050–1056.

54. Parkash, R., Rajpurohit, S. & Ramniwas, S. (2009a). Impact of darker, intermediate and lighter phenotypes of body melanization on desiccation resistance in Drosophila melanogaster. J Insect Sci, 9, 49.

55. Parkash, R., Sharma, V. & Kalra, B. (2008b). Climatic adaptations of body melanisation in Drosophila melanogaster from Western Himalayas. Fly, 2, 111–117.

56. Parkash, R., Sharma, V. & Kalra, B. (2009b). Impact of body melanisation on desiccation resistance in montane populations of *D. melanogaster*: Analysis of seasonal variation. Journal of Insect Physiology, 55, 898–908.

57. Pérochon, E., Guénard, B., Gippet, J.M.W., Klaftenberger, T., Ollier, S. & Bertelsmeier, C. (2026). Environmental conditions shape the global distribution of ant societies. Proceedings of the National Academy of Sciences, 123, e2530826123.

58. R Core Team. (2024). R: A language and environment for statistical computing. R Foundation for Statistical Computing, Vienna, Austria.

59. Rajpurohit, S., Parkash, R. & Ramniwas, S. (2008). Body melanization and its adaptive role in thermoregulation and tolerance against desiccating conditions in drosophilids. Entomological Research, 38, 49–60.

60. Ramniwas, S. & Kajla, B. (2012). Divergent strategy for adaptation to drought stress in two sibling species of *montium* species subgroup: *Drosophila kikkawai* and *Drosophila leontia*. Journal of Insect Physiology, 58, 1525–1533.

61. Ramniwas, S., Kajla, B., Dev, K. & Parkash, R. (2012). Direct and correlated responses to laboratory selection for body melanisation in *D. melanogaster* : support for melanism-desiccation resistance hypothesis. Journal of Experimental Biology, jeb.076166.

62. Rensch, B. (1929). Das Prinzip geographischer Rassenkreise und das Problem der Artbildung. Gebrüder Borntraeger, Berlin.

63. Roulin, A. (2014). Melanin-based colour polymorphism responding to climate change. Global Change Biology, 20, 3344–3350.

64. Schifani, E., Oueslati, W., Bouayed, J., Ben Aba, W., Nouira, S., Carapezza, A., et al. (2024). The mimicry complex of the acrobat ant Crematogaster scutellaris in Tunisia: Colobopsis imitans and Mimocoris rugicollis (Hymenoptera: Formicidae; Heteroptera: Miridae). Fragmenta entomologica, Vol. 56 No. 2 (2024).

65. Schultheiss, P., Nooten, S.S., Wang, R., Wong, M.K.L., Brassard, F. & Guénard, B. (2022). The abundance, biomass, and distribution of ants on Earth. Proceedings of the National Academy of Sciences, 119, e2201550119.

66. Schultz, S., Doerschner, K. & Maloney, L.T. (2006). Color constancy and hue scaling. Journal of Vision, 6, 10.

67. Skaldina, O., Peräniemi, S. & Sorvari, J. (2018). Ants and their nests as indicators for industrial heavy metal contamination. Environmental Pollution, 240, 574–581.

68. Skaldina, O. & Sorvari, J. (2017a). Not simply red: Colouration of red wood ant Formica rufa (Hymenoptera: Formicidae) is polymorphic, modular and size-dependent. Eur. J. Entomol., 114, 317–324.

69. Skaldina, O. & Sorvari, J. (2017b). Wood ant colouration as an ecological indicator for the level of disturbance in managed coniferous forests. Ecological Indicators, 72, 444–451.

70. Stuart-Fox, D., Newton, E. & Clusella-Trullas, S. (2017). Thermal consequences of colour and near-infrared reflectance. Philos Trans R Soc Lond B Biol Sci, 372, 20160345.

71. Tian, L. & Benton, M.J. (2020). Predicting biotic responses to future climate warming with classic ecogeographic rules. Current Biology, 30, R744–R749.

72. Toscani, M., Valsecchi, M. & Gegenfurtner, K.R. (2017). Lightness perception for matte and glossy complex shapes. Vision Research, 131, 82–95.

73. Vynne, C., Gosling, J., Maney, C., Dinerstein, E., Lee, A.T.L., Burgess, N.D., et al. (2022). An ecoregion-based approach to restoring the world’s intact large mammal assemblages. Ecography, 2022, ecog.06098.

74. Wagner, H.C. & Csősz, S. (2025). The “Chameleon Ant” Colobopsis imitans Adapts Its Mimetic Appearance to Local Model Species Across the Mediterranean Basin (Hymenoptera: Formicidae). Ecology and Evolution, 15, e72674.

75. Wagner, H.C., Kraker, F., Bračko, G. & Schifani, E. (2025). Extensive Field Observations Throw Light on the Evolution of Mimicry in Camponotus lateralis (Hymenoptera: Formicidae). Ecology and Evolution, 15, e72530.

76. Webb, C.O., Ackerly, D.D., McPeek, M.A. & Donoghue, M.J. (2002). Phylogenies and Community Ecology. Annual Review of Ecology, Evolution, and Systematics, 33, 475–505.

77. Wittkopp, P.J. & Beldade, P. (2009). Development and evolution of insect pigmentation: Genetic mechanisms and the potential consequences of pleiotropy. *Seminars in Cell & Developmental Biology*, A Special Edition on Biosensors and Development of Pigment Cells and Pigment Patterns, 20, 65–71.

78. Yu, D., Liu, Y., Shi, P. & Wu, J. (2019). Projecting impacts of climate change on global terrestrial ecoregions. Ecological Indicators, 103, 114–123.

